# Building the cytokinetic contractile ring in an early embryo: initiation as clusters of myosin II, anillin and septin, and visualization of a septin filament network

**DOI:** 10.1101/2021.05.25.445582

**Authors:** Chelsea Garno, Zoe H. Irons, Courtney M. Gamache, Xufeng Wu, Charles B. Shuster, John H. Henson

## Abstract

The cytokinetic contractile ring (CR) was first described some 50 years ago, however our understanding of the assembly and structure of the animal cell CR remains incomplete. We recently reported that mature CRs in sea urchin embryos contain myosin II mini-filaments organized into aligned concatenated arrays, and that in early CRs myosin II formed discrete clusters that transformed into the linearized structure over time. The present study extends our previous work by addressing the hypothesis that these myosin II clusters also contain the crucial scaffolding proteins anillin and septin, known to help link actin, myosin II, RhoA, and the membrane during cytokinesis. Super-resolution imaging of cortices from dividing embryos indicates that within each cluster, anillin and septin2 occupy a centralized position relative to the myosin II mini-filaments. As CR formation progresses, the myosin II, septin and anillin containing clusters enlarge and coalesce into patchy and faintly linear patterns. Our super-resolution images provide the initial visualization of anillin and septin nanostructure within an animal cell CR, including evidence of a septin filament network. Furthermore, Latrunculin-treated embryos indicated that the localization of septin or anillin to the myosin II clusters in the early CR was not dependent on actin filaments. These results highlight the structural progression of the CR in sea urchin embryos from an array of clusters to a linearized purse string, the association of anillin and septin with this process, and provide, for the first time, the visualization of septin filament higher order structure in an animal cell CR.

## Introduction

The mediation of the process of cytokinesis is arguably the most essential function of the actomyosin cytoskeleton in animal cells. Despite significant research efforts extending over decades, key mechanisms underlying the formation of the cytokinetic contractile ring (CR) remain poorly understood (see recent reviews by Pollard, 2017; Glotzer, 2017; Mangione and Gould, 2019; Pollard and O’Shaughnessy, 2019). This is particularly the case in animal cells, whereas in fission and budding yeast the roles of various CR-associated proteins and their structures, interactions and mechanisms have been more extensively characterized, imaged and modeled (see recent reviews by Willet et al., 2015; Meitinger and Palani, 2016; Glotzer, 2017; Marquardt et al., 2019).

Our knowledge of the CR traces back to early transmission electron microscopy (TEM) based studies performed by Schroeder (Schroeder, 1970; 1972; 1973) and others (Perry et al., 1971; Sanger and Sanger, 1980; Maupin and Pollard, 1986) that established that cytokinesis in animal cells was mediated by a circumferential ring of actin and putative non-muscle myosin II filaments. Many of these CR studies hypothesized that the organization of actin and myosin II facilitated a sliding filament-based “purse string contraction” mechanism for ring constriction. However, clear evidence of the exact architecture of the CR was lacking in these earlier works and only more recent studies employing super-resolution microscopy and head and tail-based labeling of myosin II filaments (Beach et al., 2014, Fenix et al. 2016; Henson et al., 2017) have demonstrated that myosin II within the mature CR is organized into aligned arrays that are oriented appropriately for a purse string contraction mechanism. Our previous work also extended this to the TEM level in which we used platinum replicas of cortices isolated from dividing sea urchin embryos to show a purse-string consistent orientation of both actin and myosin II filaments (Henson et al., 2017).

This recent success in defining the actin and myosin II organization in the mature animal cell CR does not discount the fact that many unanswered questions remain. For example, little is known about the pre-CR structure in animal cells, although our recent work (Henson et al., 2017) and other studies (Maupin et al. 1994; Straight et al., 2003; Werner et al., 2007, Foe and von Dassow, 2008; Zhou and Wang, 2008; Osório et al., 2019) suggest that the precursor of the CR consists of an array of myosin II-containing clusters. In fission yeast, nodes of myosin II (Myo2) contribute to the CR assembly (Pollard and Wu, 2010; Lee et al., 2012; Laplante et al., 2016) through a search-capture and pull mechanism (Vavylonis et al., 2008) and the interesting possibility exists that the clusters present in animal cells correspond to evolutionarily derived structures (Cheffings et al., 2016). However, there are a number of fundamental differences between the cell division mechanisms of yeast and those in animal cells, including yeast’s intranuclear karyokinesis, the spatial regulation of septation, and the added complication of a cell wall during cytokinesis. Therefore, despite the similarities between our actomyosin clusters and yeast nodes, we hypothesize that the CR assembly in large embryonic cells derives from a hybrid mechanism between yeast node congression and actomyosin contraction-based organization of the linear arrays of actin and myosin II filaments in the mature CR.

Questions about animal cell CR organization extend well beyond the actomyosin structure to the architecture and dynamics of two major CR scaffold proteins: septin and anillin. In budding yeast, the formation of a ring of septin filaments is a crucial step in CR assembly (Ong et al. 2014; Marquardt et al., 2019) and septin is associated with the CR in animal cells where it is thought to serve as a potential scaffold between the membrane, anillin, myosin II, and actin (Bridges and Gladfelter, 2015; Spiliotis, 2018). However, no study to date has definitively demonstrated the higher order structural organization of septins in an animal cell CR. In budding yeast, the ultrastructural arrangement of septin filaments in the bud neck has been visualized (Ong et al., 2014), although the progressive ring, hourglass and then double ring septin filament structures present are thought to be separate from the actomyosin ring (Ong et al. 2014; Marquardt et al., 2019).

Another scaffold protein critical for cytokinesis is anillin (Field and Alberts, 1995), which has been demonstrated to serve as an integrating link between the major regulator of cytokinesis RhoA and the CR components actin, myosin II, septin and formin in animal and yeast cells (Oegema et al., 2000; Maddox et al., 2005; Straight et al., 2005; Maddox et al., 2007; Hickson and O’Farrell, 2008; Piekny and Glotzer, 2008; Piekny and Maddox, 2010; Glotzer, 2017; Carim et al., 2020). In fission yeast its analogue is the protein Mid1 which is essential for the initiation of the nodes which assemble into the CR (Glotzer, 2017). However, similar to the case with septin filaments, little is known about the nanostructural organization of anillin in the animal cell CR. This fundamental uncertainty about the precise structure, function and dynamics of the two major cytokinetic scaffolding proteins argues that further work is needed to understand the complex organization of the animal cell CR (Cheffings et al., 2016; Pollard, 2017; Mangione and Gould, 2019).

Echinoderm embryos have long served as a crucial experimental model for cytokinesis research and have been used to demonstrate the existence of the CR (Schroeder, 1972), the essential role for myosin II in CR contraction (Mabuchi and Okuno, 1977), and the involvement of RhoA in the regulation of cytokinesis (Mabuchi et al., 1993; Bement et al., 2005). The sea urchin embryo also affords an approach crucial to the investigation of the 3D arrangement of ring constituents where CRs of early embryos can be isolated by adhering dividing embryos to coverslips, and then applying a stream of buffer (Henson et al., 2019). The CRs in these isolated cortical preparations from first division embryos are roughly 10X larger than CRs in cultured mammalian cells and therefore overcome the significant limitation of small cell size and membrane curvature that plague studies trying to resolve the structure of the CR in mammalian cells. The cortex isolation method also allows for visualization of the CR from the vantage point of the cytoplasmic face of the plasma membrane. Multiple investigators have used isolated sea urchin embryo cortices to examine CR structure (Yonemura and Kinoshita, 1986; Otto and Schroeder, 1990; Mabuchi, 1994; Uehara et al., 2008) including our own recent work (Henson *et al*., 2017).

In the present study we use first division sea urchin embryos to investigate the structure and dynamics of the CR components actomyosin, septin2 and anillin. We extend our previous work by testing the hypothesis that the CR scaffolding proteins anillin and septin mirror the transformation of structural organization we have reported for CR myosin II (Henson et al., 2017). We start by characterizing antibody probes for sea urchin anillin and septin2 and then investigate their localization in the CR relative to myosin II via super-resolution imaging. Our results indicate that septin2 and anillin are affiliated with myosin II in the band of clusters/nodes that appears to serve as a precursor to the CR early in the cell division process. Within each cluster, anillin and septin2 both occupy a more central position relative to the myosin II minifilament head groups. As CR formation progresses, the septin2 and anillin staining focus into a narrower pattern coincident with a similar change in activated myosin II minifilament distribution. Both 3D SIM and STED imaging provide the first clear evidence of the existence of a network of septin filaments affiliated with myosin II in mature CRs. Furthermore, the localization of either septin2 or anillin to the early CR myosin II clusters was not dependent on actin filaments given that it occurred in embryos treated with Latrunculin prior to cell division. Taken together these results underscore the structural evolution of the CR in sea urchin embryos from an array of clusters to a linearized purse string, the critical scaffolding roles that anillin and septin play based on their localizations with expected binding partners, and provide, for the first time, the visualization of higher order septin filaments in the CR of an animal cell.

## Materials and Methods

### Animals, antibodies, and reagents

*Lytechinus pictus* sea urchins were purchased from Marinus Scientific (Lakewood, CA) and *Strongylocentrotus purpuratus* sea urchins were collected from the waters surrounding Port Townsend, WA, and maintained at the Friday Harbor Laboratories (Friday Harbor, WA). All animals were kept in either running natural sea water or closed artificial sea water systems at 10-15°C.

Primary antibodies used included a rabbit polyclonal antibody raised against sea urchin egg myosin II heavy chain isolated via ATP-based precipitation of actomyosin from *S. purpuratus* egg extracts and electrophoretically purified (Henson et al., 1999), a mouse monoclonal antibody against the Ser19 phosphorylated form of myosin II regulatory light chain (P-MyoRLC) from Cell Signaling Technology (Danvers, MA), a rabbit monoclonal antibody against a peptide from human septin 2 from Abcam, Inc (Cambridge, MA), a mouse monoclonal antibody against a conserved epitope of chicken gizzard actin (clone C4) from EMD, Millipore (Burlington, MA), a rat monoclonal antibody against yeast alpha-tubulin (clone YL1/2) from Thermo Fisher Scientific (Pittsburgh, PA), and a mouse monoclonal antibody against human RhoA/B/C (clone 55) from Upstate Biotechnology (Lake Placid, NY). In order to generate a sea urchin anillin antibody, the PH domain of *S. purpuratus* anillin (aa 980-1124, Accession #XM_030975862) was cloned into pET302, expressed in *E. coli*, and the purified polypeptide used as the immunogen for a commercially prepared rabbit polyclonal antiserum (Thermo Fisher Scientific) which was then affinity purified using the anillin PH domain. Appropriate secondary antibodies conjugated to Alexa Fluor 488, 555, 568, or Oregon Green as well as Alexa Fluor 488, 633, and 647 conjugated phalloidin were obtained from Thermo Fisher Scientific. Latrunculin A and B (LatA; LatB) were obtained from Cayman Chemical (Ann Arbor MI). Unless otherwise indicated, the majority of reagents were purchased from either Sigma-Aldrich (St. Louis, MO) or Thermo Fisher Scientific.

### Gamete and coelomocyte collection, fertilization, and cleavage cortex isolation

Sea urchin gametes were collected via intracoelomic injection with 0.5 M KCl, with sperm collected dry and eggs spawned in either natural sea water or MBL artificial sea water (ASW: 423 mM NaCl, 9 mM KCl, 9.27 mM CaCl_2_, 22.94 mM MgCl_2_, 25.5 mM MgSO_4_, 2.14 mM NaHCO_3_, pH 8.0) and subsequently dejellied by multiple washing with ASW. Eggs were fertilized by addition of dilute sperm, the fertilization envelopes removed using 1 M urea (pH 8.0), and then washed into and reared in MBL calcium free sea water (MBL ASW minus CaCl_2_ and plus 1 mM EGTA) at 10-15°C. For disruption of actin filaments embryos were treated with either 1 µM LatA or 20 µM LatB starting at 30-60 min prior to first division. Cleavage cortices were generated as described in Henson et al. (2019). In brief, embryos at the appropriate stage of cell division were allowed to quickly settle onto poly-L-lysine (2 mg/ml) coated coverslips and then exposed to fluid shear force from a pipette containing an isotonic cortex isolation buffer (CIB: 0.8 M mannitol, 5 mM MgCl_2_, 10 mM EGTA, 100 mM HEPES, pH 7.4). Isolated cortices were rinsed twice in CIB prior to further processing for fluorescent staining. Sea urchin coelomocytes were isolated from the perivisceral fluid of adult animals and maintained in coelomocyte culture media (0.5 M NaCl, 5 mM MgCl_2_, 1 mM EGTA, and 20 mM HEPES, pH 7.2) as described in Smith et al. (2019).

### Fixation, fluorescent staining and microscopic imaging and analysis

Embryos – either attached to poly-L-lysine coated coverslips or kept in suspension – were fixed in Millonig’s fixative (0.2 M NaH_2_PO_4_, 0.136 M NaCl, pH 7.0) containing 3.7% formaldehyde for 20 minutes, washed 3X and then left overnight in 0.1% Triton X-100 in phosphate buffered saline (PBST), and then blocked overnight in 3% BSA diluted in PBST. Isolated cortices were fixed in 2-3% formaldehyde in CIB for 5 min followed by blocking in 2% goat serum and 1% BSA in PBS for at least 30 minutes. For immunolocalization of RhoA/B/C, cortices were fixed in 10% trichloroacetic acid following the methods of Yonemura et al. (2004). Coelomocytes were settled and fixed as described in Smith et al. (2019). Immunostaining of all samples was performed with appropriate primary and secondary antibodies diluted in blocking buffer with embryos typically being stained in each antibody step overnight whereas cortices and coelomocytes were stained for 1 hour in each stage. Fluorescent phalloidin was added to the secondary antibody staining step. Samples for conventional and 3D-SIM microscopy were typically mounted in nonhardening Vectashield antifade mounting media (Vector Laboratories, Burlingame, CA) plus or minus DAPI prior to imaging. Samples for STED imaging were mounted in Prolong Diamond mounting media (Thermo Fisher).

Conventional epifluorescence microscopy of samples was performed on a Nikon (Tokyo, Japan) 80i microscope using either a 40X/0.75 NA Plan Fluorite or 60X/1.4 NA Plan Apo phase contrast objective lens with digital images captured using a Photometrics (Tuscon, AZ) CoolSnap Cf cooled CCD camera. Confocal microscopy was performed on either an Olympus (Tokyo, Japan) Fluoview 500 laser scanning confocal microscope using a 40X/1.15 NA UApo water immersion DIC objective lens, or an Andor Dragonfly 505 spinning disk confocal system (Oxford Instruments, Abingdon, UK) using either an Olympus 60X/1.3 NA silicone or a 100X/1.49 NA oil immersion objective lens. Spinning disk confocal digital images were acquired with an Andor sCMOS EMCCD camera.

Super-resolution microscopy was performed using two different methods. For 3D structured illumination microscopy (3D-SIM, Gustafsson et al., 2008) we utilized a DeltaVision OMX 3D-SIM Imaging System (GE Healthcare Bio-Sciences, Pittsburgh, PA) with an Olympus 60X/1.42 NA objective lens. Captured images were reconstructed using SoftWoRx software. Stimulated Emission Depletion (STED) super-resolution microscopy (Hell and Wichmann, 1994) was performed on a Leica (Wetzlar, Germany) Sp8 STED confocal using a 100X/1.4 NA objective lens.

For all forms of microscopic images, processing and analysis was performed using either Fiji/ImageJ (Bethesda, MD) or Bitplane Imaris (version 8.1-9.1.2; Andor). Graphs were prepared and statistical analysis carried out using GraphPad Prism 8 (San Diego, CA), with box and whisker plots having the following attributes: the box extends from the 25^th^ to the 75^th^ percentiles, the whiskers extend to the minimum and maximum values, and the line in the middle of the box is the median. Final figures were prepared using Adobe Photoshop (San Jose, CA).

### Gel sample lysates and immunoblotting

Sea urchin egg/embryo gel sample lysates were generated by pelleting cells and then adding hot 2X sample buffer (625 mM Tris-HCl pH 6.8, 25% glycerol, 2% SDS, 0.01% bromophenol blue, 5% β-mercaptoethanol) at five times the volume of packed cells followed by boiling for 5 minutes. Gel samples were loaded into a 4-15% Mini-PROTEAN TGX precast protein gel (Bio-Rad) and transferred onto an Immobilon PDF membrane (Millipore) for probing. Blocking and antibody incubation was performed in TBS-Tween supplemented with 5% instant milk. Bound antibody detection was performed for septin2 immunoblots using alkaline phosphatase conjugated secondary antibodies and a BCIP+NBT substrate. For anillin immunoblots HRP conjugated secondary antibodies were used and detected by chemiluminescence using a Immun-Star HRP kit, with imaging performed on a ChemiDoc XRS molecular imaging system from Bio-Rad (Hercules, CA).

## Results

### Characterization of antibodies against the sea urchin anillin PH domain and septin2

Sea urchin anillin is predicted to contain the domains that confer anillin with its unique CR integration functions (Piekny and Maddox, 2010). These include a N-terminal domain (NTD) with a formin binding site (FBD), nearby myosin II and actin binding domains (MBD and ABD), an anillin homology domain (AHD) which interacts with RhoA, and a C terminus pleckstrin homology (PH) domain which allows for binding with septin and membrane phospholipids (Figure 1A). The PH domain of *S. purpuratus* anillin was cloned and used to generate a rabbit polyclonal antiserum which was affinity purified against the recombinant anillin PH domain and its specificity confirmed by immunoblotting (Figure 1B). Rabbit anti-sea urchin anillin reacted with the purified recombinant 16 kDa anillin PH domain antigen, and with a ∼120 kDa species in *S. purpuratus* egg and embryo lysates consistent with the predicted molecular mass of *S. purpuratus* anillin (Figure 1B).

**Figure 1.**
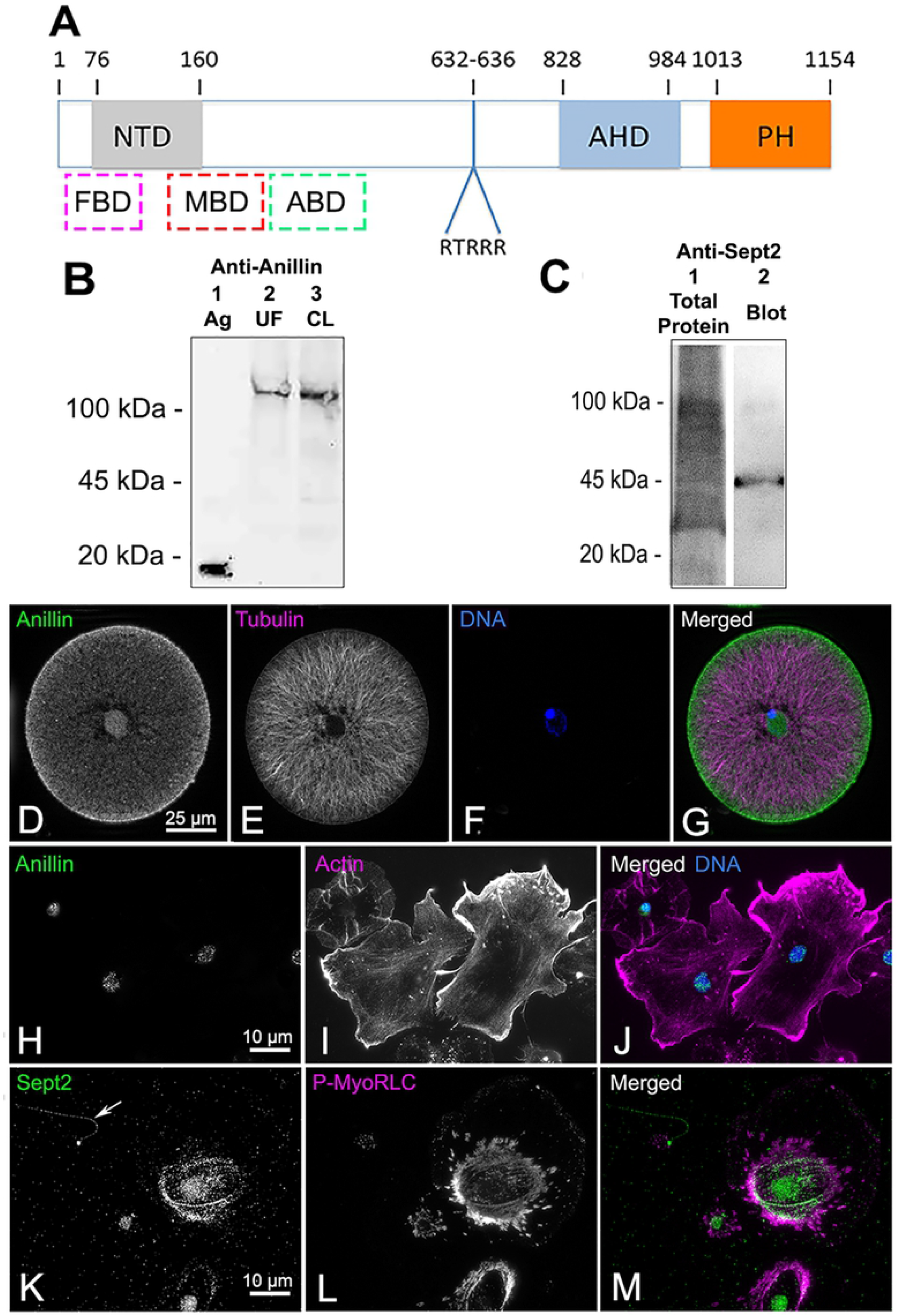
Characterization of antibodies against sea urchin anillin and septin2. (A) Starting at the N terminus, the predicted domains of sea urchin anillin include a formin binding domain (FBD), a N-terminal domain (NTD), a myosin II binding domain (MBD), an actin binding domain (ABD), a candidate nuclear localization sequence (RTRRR), an anillin homology domain (AHD), and a plecstrin homology domain (PH). The dashed boxes for FBD, MBD, and ABD indicate approximate locations. (B) Immunoblot of affinity purified anti-sea urchin anillin PH domain antibody against the purified peptide antigen (Ag, lane 1) shows the expected 16 kDa immunoreactive band. Blotting this antibody against lysates of either unfertilized *S. purpuratus* eggs (UF, lane 2), or first cleavage embryos (CL, lane 3) reveals a ∼120 kDa immunoreactive band. (C) Immunoblot of anti-human septin2 peptide antibody against *L. pictus* first cleavage embryo lysate (lane 1 = total protein; lane 2 = immunoblot) reveals a ∼45 kDa immunoreactive species. (D-J) Anillin (green) staining of a *S. purpuratus* sperm aster stage early embryo (D-G) and adult coelomocytes (H-J) shows that anillin localizes to the nucleus and in early embryos to the cortical region. Embryos are co-labeled for microtubules (magenta) and DNA (blue), whereas coelomocytes are co-labeled for actin filaments (magenta) and DNA (blue). (K-M) Septin2 (green) staining of adult *S. purpuratus* coelomocytes double labeled for P-MyoRLC (magenta) demonstrates the expected staining of septin in stress fiber-like actomyosin bundles in phagocytes and in the flagella of vibratile cells (arrow in K).

Septins are a family of GTP-binding proteins that form filaments composed of linear arrays of hetero-oligomers of different isoforms which vary across species (Bridges and Gladfelter, 2015), and the sea urchin *S. purpuratus* genome has been shown to encode homologues of the human Sept2, Sept3, Sept6, and Sept7 subgroups (Cao et al., 2007). Immunoblotting of an *L. pictus* embryo lysate with a commercial rabbit monoclonal antibody against a peptide antigen (between aa 250-350) from human Sept2 resulted in an immunoreactive band of ∼45 kDa which is similar to the 42 kDa molecular mass of human Sept2 (Figure 1C).

The sea urchin anillin and septin2 antibodies were initially further characterized by testing for expected immunofluorescence staining patterns. Anillin proteins have a nuclear localization sequence (NLS) and have been shown to be localized in the nucleus during interphase in some species (Field and Alberts, 1995; Oegema et al., 2000; Kim et al., 2017). Sea urchin anillin has a weak putative NLS (aa 632-636, Figure 1A) and staining of newly fertilized, sperm aster stage embryos shortly following syngamy with the anti-sea urchin anillin antibody showed it was present within the zygote nucleus as well as in the cortex (Figure 1D-G). Nuclear localization of anillin in interphase cells was also observed in isolated adult sea urchin coelomocytes that are terminally differentiated, post-mitotic cells (Figure 1H-J). Septin filaments are known to localize in stress fibers in cultured cells (Spiliotis, 2018) and in cilia and flagella (Palander et al., 2017). Staining adult coelomocytes with the septin2 antibody showed clear localization in the stress fiber-like actomyosin bundles of large phagocytes and in the flagellar axonemes of vibratile cells (Figure 1K-M).

### Anillin and septin2 localize to the CR of whole embryos and isolated cleavage cortices

During first division of sea urchin embryos, overlapping microtubules from the asters deliver the centralspindlin/RhoGEF complex to the equator that activates RhoA (Bement et al., 2005; Argiros et al., 2012; Su et al., 2014), which in turn is thought to help recruit anillin and activate CR actomyosin contraction. Dividing embryos stained for microtubules and anillin showed that the equatorial cortex accumulation of anillin – which was not apparent in late anaphase (Figure 2A-C) – appeared coincident with the invagination of the cleavage furrow in early telophase (Figure 2D-F), as has been described in other cell types (Field and Alberts, 1995; Oegema et al., 2000; Hickson and O’Farrell, 2008; Kim et al., 2017). As the cleavage furrow ingressed with the progression of telophase, anillin staining became highly concentrated in the region of the cleavage furrow (Figure 2D-O) - with off axis images showing a clear ring shaped anillin-stained structure (Figure 2J-L) - and remained associated with the midbody formation at the end of cytokinesis (Figure 2M-O).

**Figure 2.**
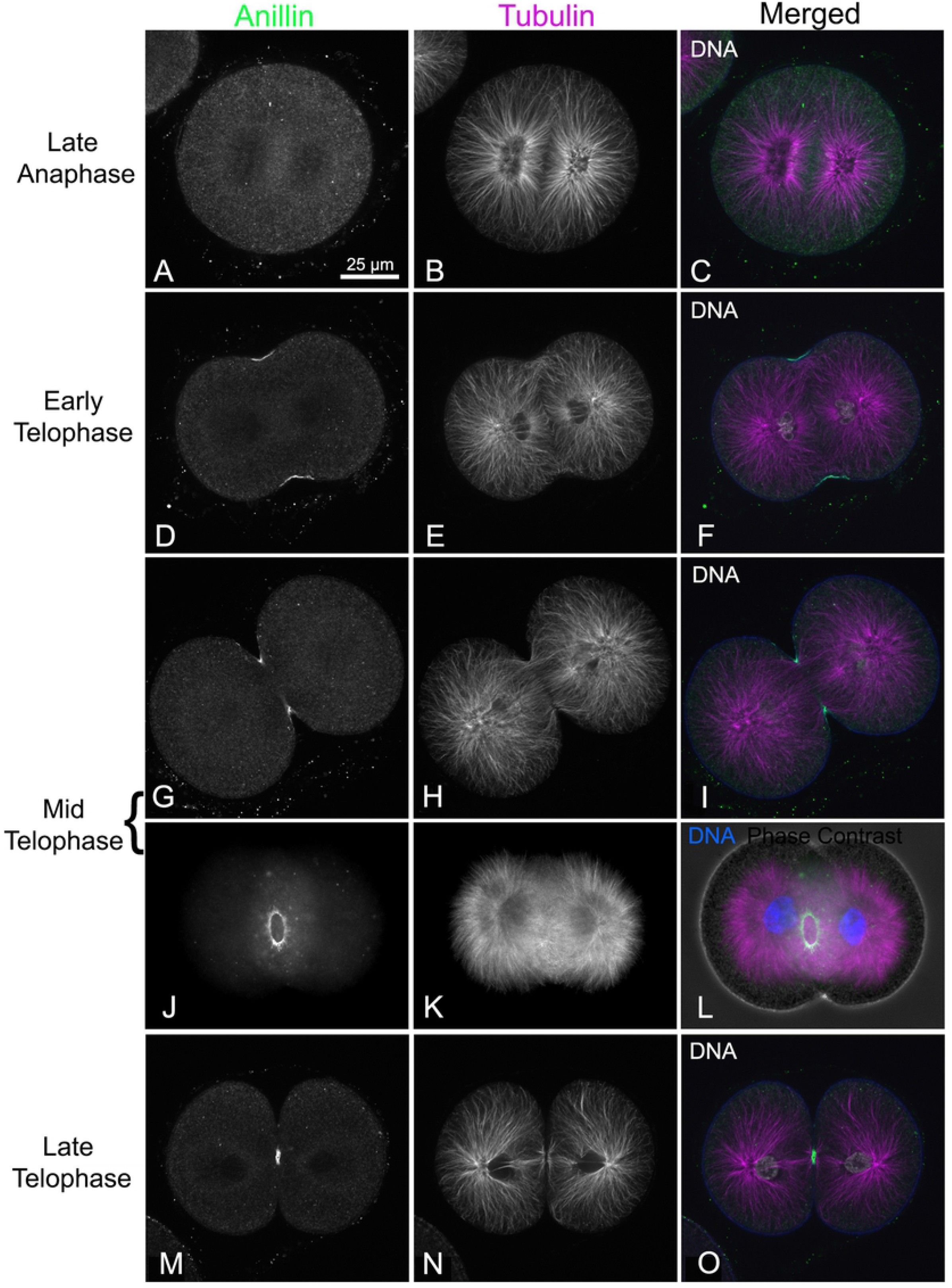
Anillin localizes to the cleavage furrow in first division embryos. In dividing *S. purpuratus* embryos co-labeled for microtubules (magenta) and DNA (white; blue in L) in order to determine mitotic progression, anillin (green) staining is not obvious in late anaphase (A-C) but begins concentrating in the early cleavage furrow starting at the initiation of telophase (D-F) and continues through the midbody stage prior to abscission in late telophase (G-O). In off axis images the anillin staining appears as an entire ring similar to other CR markers (J-L). In L the phase contrast image is superimposed on the fluorescence image for context. The magnifications of A-O are equivalent and A-I plus M-O are confocal images whereas J-L are widefield images.

The CR in fixed sea urchin embryos can be readily stained using antibodies against the Ser19 phosphorylated form of the myosin II regulatory light chain (P-MyoRLC) which is indicative of activated myosin II that is capable of actin-based contraction (Foe and von Dassow, 2008; Argiros et al., 2012; Henson et al., 2017). Localization of either septin2 or anillin with P-MyoRLC in whole embryos showed a clear association with the CR that began as a broad band of clustered, reticular staining (Figure 3A-D, M-P) which then concentrated into a narrow ring as furrowing progressed (Figure 3E-H, Q-T), and culminated in retention in the midbody (Figure 3I-L).

**Figure 3.**
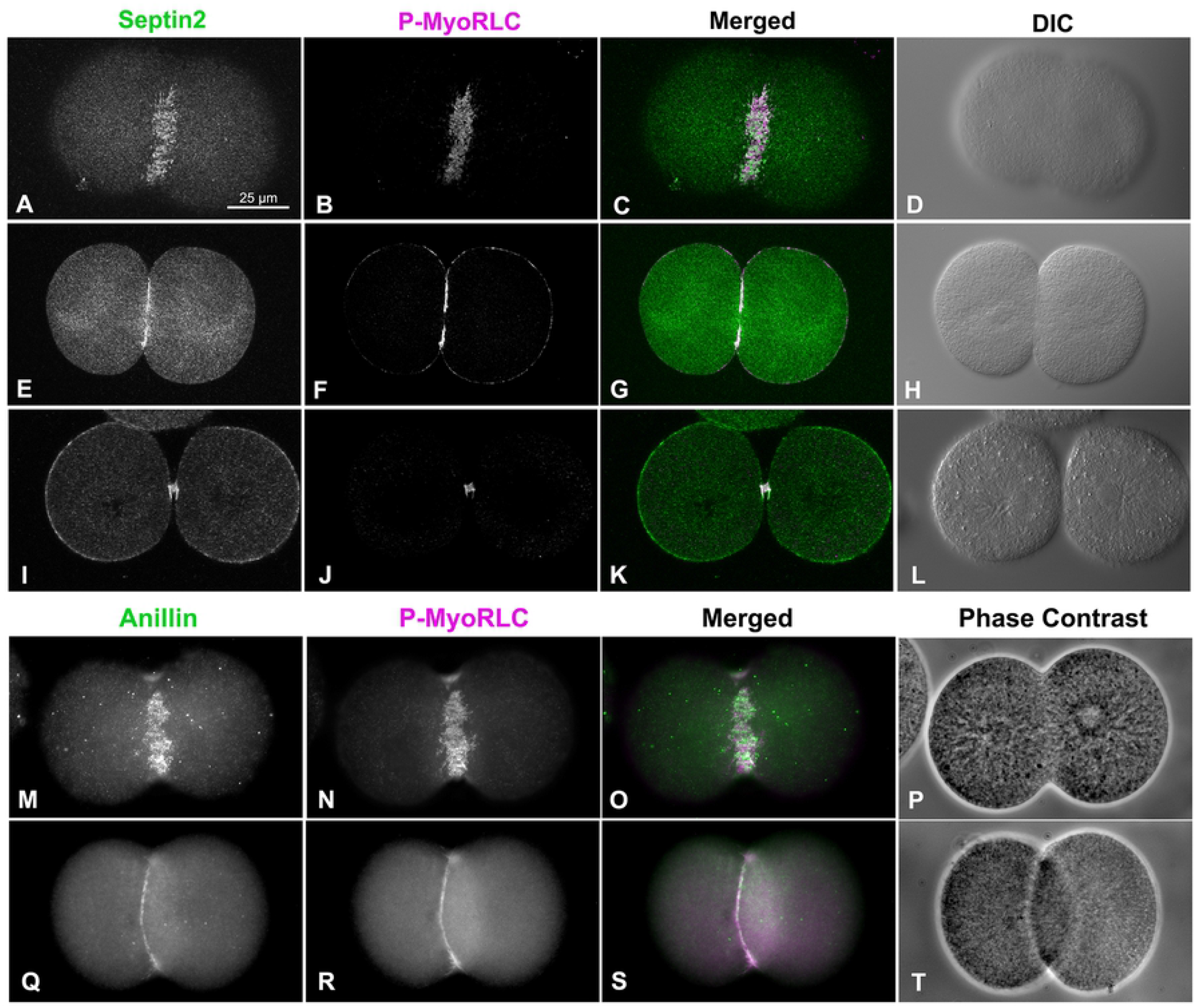
Sept2 and anillin colocalize with the CR marker P-MyoRLC in first division embryos. Both sept2 (A-L, green) and anillin (M-T, green) mirror P-MyoRLC (A-T, magenta) staining in dividing embryos and begin as collections of clusters (A-D, M-P) that progress to tight rings (E-H, Q-T), and end concentrated in the midbody (I-L). *L. pictus* embryos appear in confocal images in A-L, *S. purpuratus* embryos appear in widefield images in M-T, and the magnifications of A-T are equivalent.

Septin2 and anillin also localized with P-MyoRLC in the CR regions of cortices isolated from dividing embryos (Figure 4; 5). In examining cortices from a time series of isolations over the course of cytokinesis, those from earlier time points tended to display a broad band of punctate clusters or nodes which contained septin2, anillin and P-MyoRLC (Figure 4A-D, I-L; 5A-D). Widefield imaging of these clusters suggested an incomplete overlap between the localization of septin2 or anillin staining and that of P-MyoRLC. In cortices isolated in the mid to late stages of cytokinesis, the septin2, anillin and P-MyoRLC staining patterns (Figure 4E-H, M-Q) all appeared to have coalesced into a patchy and more linearized pattern that became very concentrated in the latest stage cortices (Figure 5E-L). Quantification of the myosin II staining patterns of 247 cortices from 3 separate experiments showed that in cortices isolated early in cytokinesis (Figure 4R) punctate CR staining predominated (70% of cortices with CRs; Figure 4T), whereas in cortices isolated in mid-late cytokinesis (Figure 4S) patchy/linear CR staining patterns were more prevalent (76% of total cortices with CRs; Figure 4T). Localization of filamentous actin in cortices using phalloidin indicated that it codistributed with the clusters in early stages (Figure 5A-D) and the condensed CR patterns present in late stages (Figure 5E-L), as well as staining the microvilli present in these cortices (Figure 5C,G,K). RhoA/B/C was also enriched in the CR region containing anillin (Figure 5M-P), septin2, actin and myosin II.

**Figure 4.**
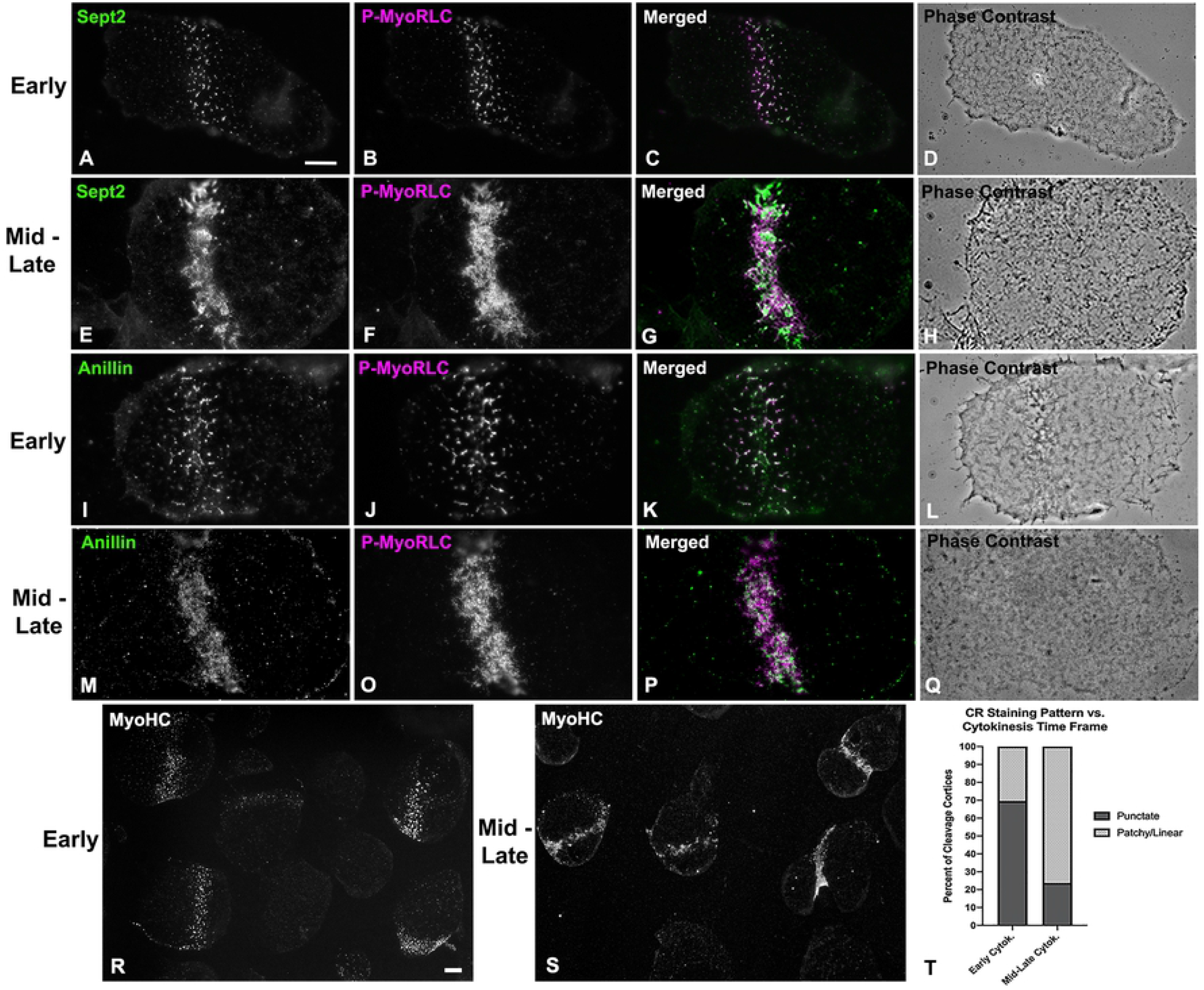
Widefield imaging of isolated cortices shows that septin2 and anillin co-distribute with active myosin II and progress from clusters to more linearized arrays. Sept2, and active myosin II (P-MyoRLC) localize together in the CR regions of isolated cortices (A-H) and progress from regularly spaced clusters in early stages (A-D) to dense assemblages of more patchy and linear structures in late stages (E-H). Anillin displays a similar co-distribution with active myosin II, as well as the analogous evolution from clusters in early cortices (I-L) to denser and more filamentous arrays in mid-late cortices (M-Q). Lower magnification images of myosin II (MyoHC) staining in cortices isolated early in cytokinesis (R) shows the presence of punctate clusters in a majority of cortices containing CRs (percentages graphed in T), whereas cortices isolated mid-late in cytokinesis (S) have a majority of cortices with patchy/filamentous CR patterns (percentages graphed in T). Bar in A = 10 µm and magnifications of A-Q are equivalent. Bar in R = 10 µm and magnification of R and S are equivalent. All cortices from *S. purpuratus* embryos.

**Figure 5.**
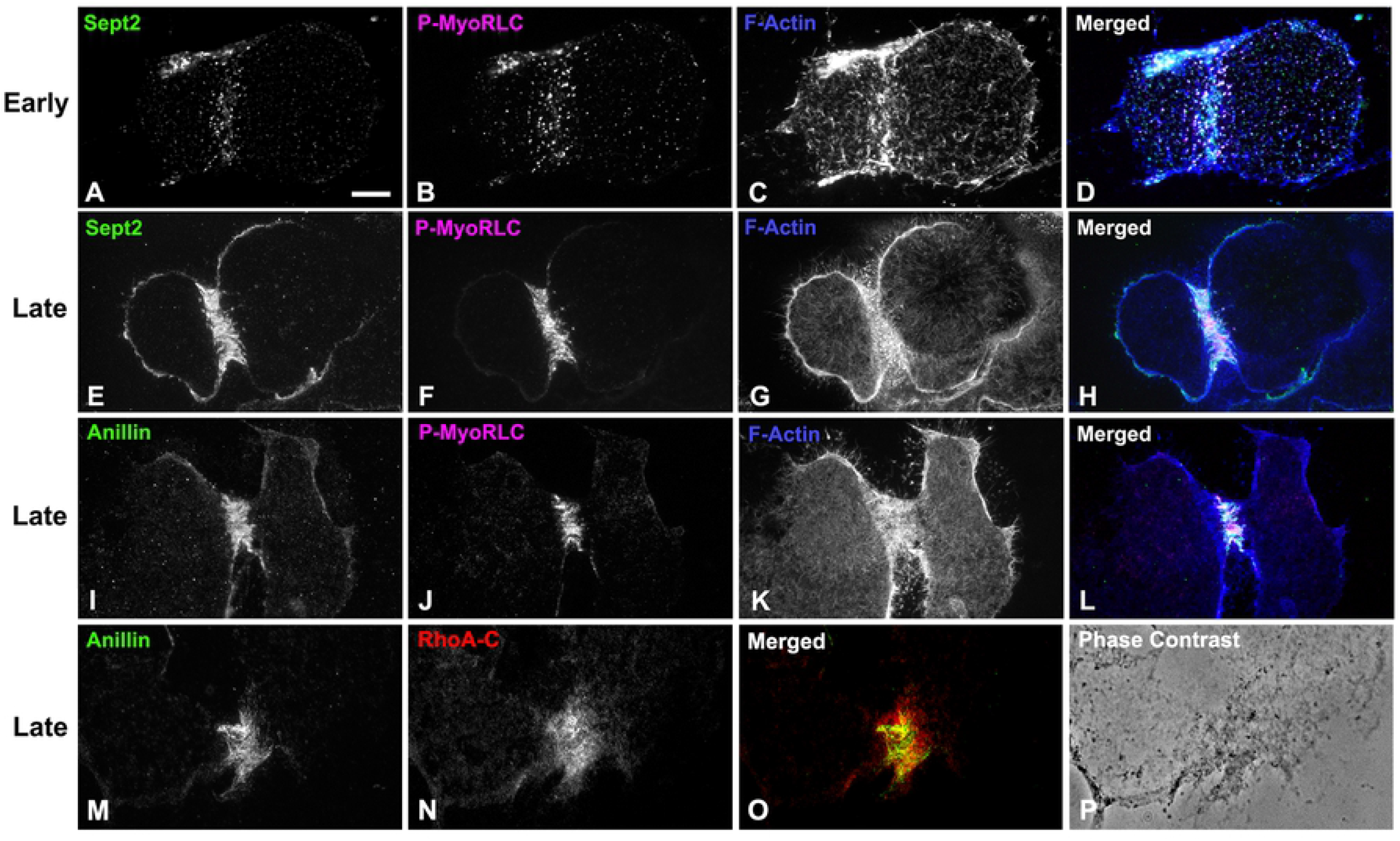
Widefield imaging of isolated cortices demonstrates that septin2 and anillin colocalize with active myosin II, filamentous actin and RhoA/B/C in CRs. Septin2, active myosin II (P-MyoRLC) and F-actin staining all associate with clusters in early CRs (A-D), and with the denser, more linear arrays in late stage CRs (E-H). Anillin displays a similar association with active myosin II and F-actin in late stage CRs (I-L) and also codistributes with Rho A/B/C in the CR (M-P). Bar in A = 10 µm; magnifications of A-L are equivalent. All cortices from *S. purpuratus* embryos.

### Super-resolution microscopy reveals the organization of myosin II, anillin and septin2 clusters that serve as CR precursors

Consistent with our widefield imaging (Figure 4; 5), 3D-SIM imaging of early cleavage stage cortices demonstrated that myosin II, septin2 and anillin are all organized in a broad band of regularly spaced clusters (Figure 6). Within these ring or stellate shaped clusters, the distribution of the myosin II heavy chain (MyoHC) tail and the P-MyoRLC head antibody probes (characterized in Henson et al., 2017) often indicated an extensive overlap of these two regions, suggesting that myosin II filaments may be densely packed (Figure 6A,D,G). The myosin II stained clusters would often form rings with hollow centers (Figure 6D; 7B,C) in which it appeared that myosin II minifilaments were arrayed around the periphery in a head to head circular structure (Figure 7B,C). In some stellate-shaped clusters the myosin II minifilaments had adopted the linear alignment of head-to-head chains or networks (Figure 6D,G; 7A) that our previous TEM images have shown to exist in the CR in embryos and in the cytoskeleton of coelomocytes (Henson et al., 2017). Z axial images indicated a significant superposition between the two myosin II probes, with the suggestion that the myosin II heads might be farther away from the membrane (bottom of image) than the tail regions (Figure 6J). However, the lower resolution and distorted nature of 3D-SIM Z axial images make it difficult to draw any definitive conclusions.

**Figure 6.**
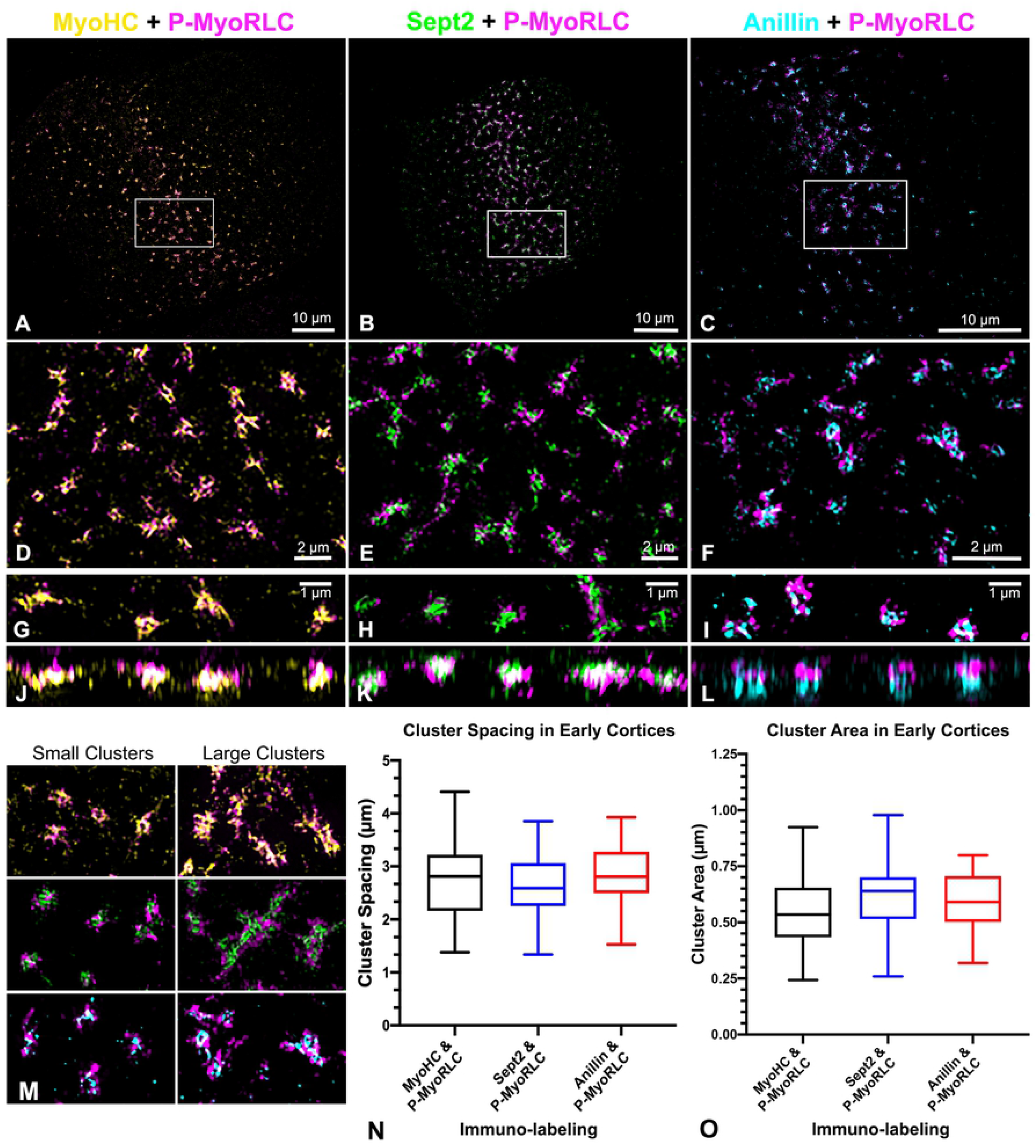
3D-SIM super-resolution imaging of clusters of myosin II, septin2 and anillin in early CR stage cortices. (A-F) Survey (A-C) and higher magnification views (D-F: enlarged white boxes in A-C) of isolated cortices from dividing embryos double labeled for P-MyoRLC (magenta in A-F) and either MyoHC (yellow in A,D), septin2 (green in B,E), or anillin (cyan in C,F). (G-L) The pairs of images that appear in G&J, H&K and I&L consist of a 10 µm x 3 µm XY image on the top paired with a corresponding 10 µm x 2 µm XZ image of the same clusters on the bottom. (M) In later stage CR regions of isolated cortices clusters become enlarged and appear to interact/coalesce with one another. (N,O) Box and whisker plots (min/max with line at median) of cluster spacing and cluster area in early stage cortices stained by the three combinations of antibodies. Bar magnifications as indicated, with images in M equivalent in magnification to panel F. All cortices from *S. purpuratus* embryos.

**Figure 7.**
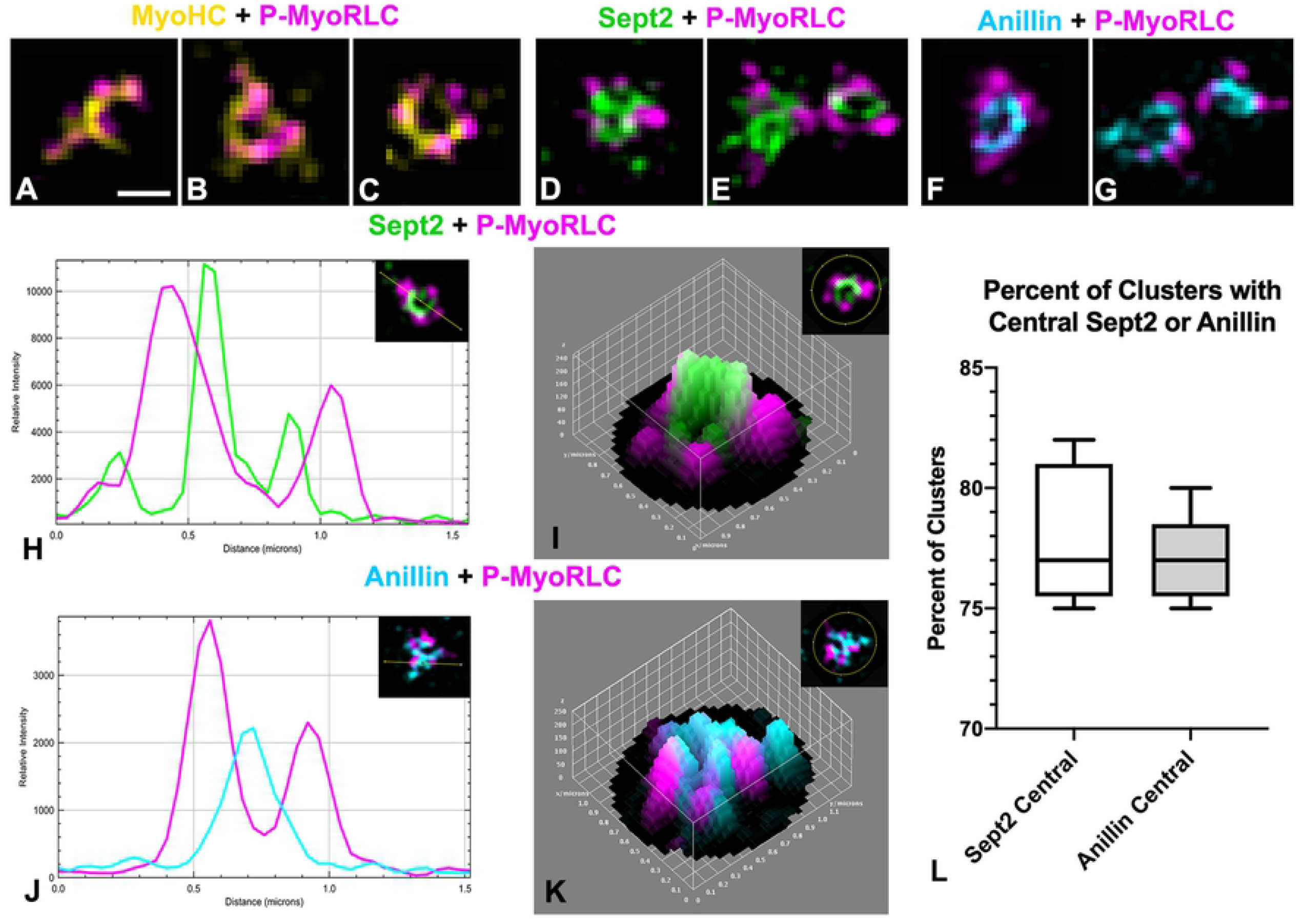
Analysis of cluster organization using 3D SIM shows that septin2 and anillin tend to be central with myosin II on the periphery. (A-C) MyoHC (yellow) and P-MyoRLC (magenta) staining of early cytokinesis stage clusters showing what appear to be mini-filaments arranged in chains (A) and rings (B,C). (D-G) Staining of P-MyoRLC (magenta) with either septin2 (green in D,E) or anillin (cyan in F,G) shows peripheral position of myosin II heads and the more central position of septin2 and anillin. (H-K) The central location of septin2 (green in H,I) and anillin (cyan in J,K) was confirmed by analyzing clusters with 2D line scans (H,J) and 3D surface plots (I,K) of relative staining intensities. Insets in H-K show images being analyzed and the line or area ROI – the images of clusters in I and K have been rotated to match the orientation of the 3D surface plots. (L) Box and whisker plots (min/max with line at median) of the quantitation of the percent of total clusters with centralized septin2 or anillin in early stage cortices. Bar in A = 500 nm; magnifications of A-G are equivalent. *S. purpuratus* cortices in A-G.

Imaging of septin2 and P-MyoRLC stained clusters in cortices with 3D-SIM (Figure 6B,E,H; 7D,E) indicated that septin2 often appeared to be in the center of clusters relative to the myosin II head staining and this was corroborated by 2D line scans and 3D surface plots of the relative intensity of sept2 and P-MyoRLC staining in clusters (Figure 7H,I) – with an average of 78% of clusters containing centralized septin2 (Figure 7L). The pattern of septin2 labeling was often ring-shaped and in some cases resembled short segments of filaments (Figure 6E,H). The Z axial images of septin2 and P-MyoRLC (Figure 6K) showed overlap and no clear distinction in the orientation of the two labels. 3D-SIM imaging of anillin and P-MyoRLC demonstrated that anillin also appeared to occupy the cluster centers and was organized in punctate, ring or C-shaped structures that appeared in close proximity to the myosin II head probe (Figure 6C,F,I; 7F,G). 2D line scans and 3D surface plots of the anillin and P-MyoRLC relative intensities (Figure 7J,K) corroborated anillin’s central location, with 77% of clusters analyzed showing centralized anillin (Figure 7L). Interestingly, the Z axial images of anillin and P-MyoRLC (Figure 6L) showed a clear distinction in the orientation of the anillin and myosin II head probes with the anillin being closer to the membrane (bottom of image in Figure 6L). For quantification of septin2 and anillin positioning in clusters, 299 total clusters were analyzed from three separate experiments.

Examination of cortices isolated in a time series showed that clusters from cortices later in cytokinesis became larger in size and more extensive (Figure 6M) although they appeared to retain the overall organization of myosin II, septin2, and anillin apparent in smaller clusters. Taken together these images suggest that these myosin II, septin2 and anillin clusters are precursors of the CR and that their growth and coalescence over time generates the mature structure of the CR. They also indicate a codependence in terms of localization given that clusters consistently showed the presence of myosin II and either septin2 or anillin. In terms of physical parameters, the small clusters in early cortices were spaced an average of 2.7 µm apart (Figure 6N) and averaged 586 nm^2^ in area (Figure 6O), with these measurements not being significantly different in clusters stained with any of the three antibody combinations. This lack of difference is important because it provides assurance that the clusters identified with the three different antibody combinations are the same structures. In terms of comparing the physical parameters of the sea urchin embryo clusters to the nodes that initiate cytokinesis in fission yeast, the sea urchin clusters are larger and more widely spaced as would be expected given the large difference in size between the two cells. In fission yeast nodes are ∼500-700 nm apart and average ∼100-200 nm in diameter (Wu et al., 2006; Laporte et al., 2011; Laplante et al., 2016).

### Super-resolution microscopy demonstrates the organization of myosin II, anillin and septin2 in the mature CR and provides evidence of a septin filament network

In isolated cortices containing mature CRs 3D-SIM imaging demonstrated that the pattern of myosin II labeling was linearized with an alignment of chains of head-to-head associated myosin II filaments (Figure 8A-D), as we have previously reported (Henson et al., 2017). In contrast, septin2 staining was organized into a honeycomb pattern network that was closely associated with P-MyoRLC staining (Figure 8E-H). The septin2 staining within the network was frequently discontinuous/periodic which may be the result of the septin2 antibody binding to only one of the septin isoforms within the linear structure of a heteropolymer. The overall network pattern of septin2 within the CR appeared similar to the gauze-like organization of septin filaments that has been reported *in vitro* (Bertin et al., 2010; Garcia et al., 2011) and *in vivo* in other cell types (Rodal et al., 2005; Bertin et al., 2012; Dolat et al., 2014). One criticism of 3D-SIM microscopy is that the structured illumination itself and/or the associated post-acquisition processing can create a honeycomb pattern artifact in the images (Demmerle et al., 2017). Therefore, in order to corroborate the septin2 network staining seen using 3D-SIM, we imaged cortices with STED microscopy, a higher resolution form of super-resolution microscopy (Schermelleh et al., 2010) that does not suffer from this type of artifact. STED images of cortices also showed clear examples of a septin2 filament network in the CR intimately affiliated with myosin II (Figure 8I-L; 9A-C). We believe that our 3D-SIM and STED images provide the first definitive visualization of septin filament higher order structure in the CR of an animal cell.

**Figure 8.**
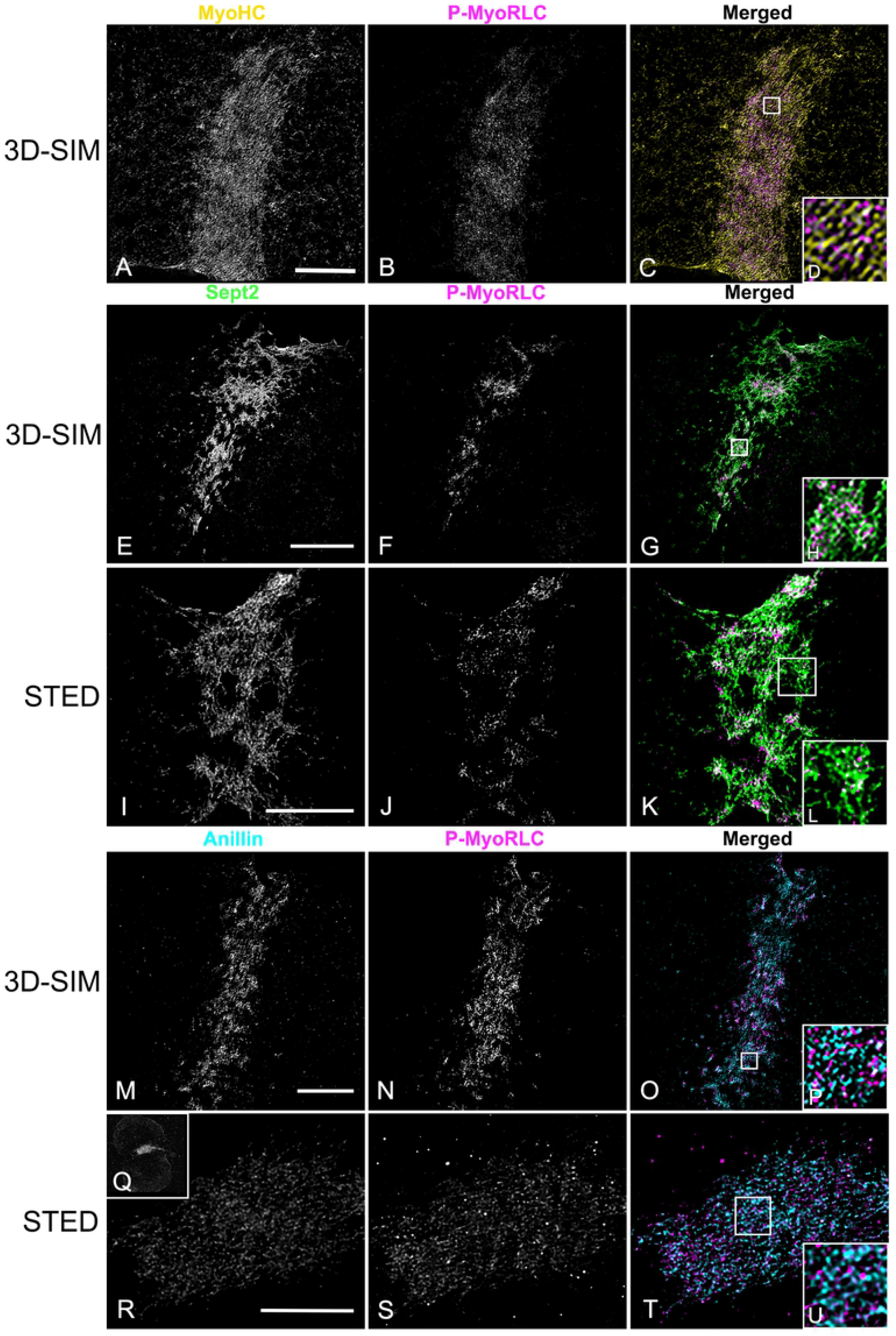
3D-SIM and STED super-resolution imaging of myosin II, septin2 and anillin in late stage CRs in isolated cortices. (A-D) MyoHC (yellow) and P-MyoRLC (magenta) staining of mature CR showing alignment of head-to-head minifilament chains (C,D). (E-L) Septin2 (green) and P-MyoRLC (magenta) labeling of a late stage CR region showing the network-like structure of septin2 filaments in close association with myosin II (H,L) using both 3D-SIM (E-H) and STED (I-L) imaging. (M-U) Anillin (cyan) and P-MyoRLC (magenta) staining of a mature CR indicates that anillin is more punctate in distribution and in close proximity to myosin II (M-P). The cortex in Q-V is highly contracted (Q shows a low magnification confocal view) and in this CR remnant the STED imaging of anillin appears similar to a network. White boxes in C,G,K,O,T correspond to the regions that appear at higher magnification in the insets labeled D,H,L,P,U. Bars = 10 µm. All cortices from *S. purpuratus* embryos.

**Figure 9.**
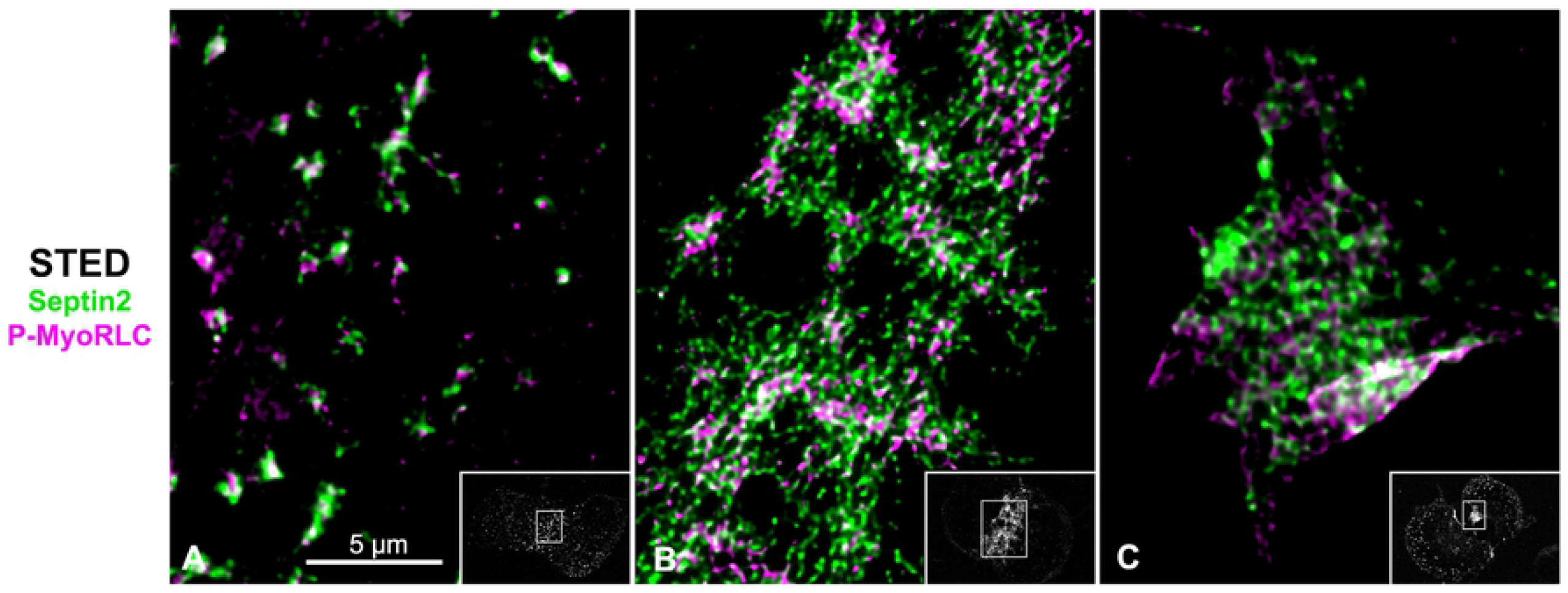
STED imaging of CR septin2 in isolated cortices shows a transformation from clusters to a filamentous network. (A-C) STED imaging of septin2 (green) and P-MyoRLC (magenta) in isolated *S. purpuratus* cortices shows elongate septin2 filaments in enlarging clusters in an early CR (A) versus the filamentous network structure of septin2 staining in later stage CRs (B,C). Insets in A-C show lower magnification confocal images of the cortices with the regions imaged by STED outlined by a white rectangle. Bar = 5 µm; magnifications of A-C are equivalent.

Imaging of anillin stained mature stage CRs in cortices with 3D-SIM (Figure 8M-P) showed that unlike septin2, the anillin pattern often appeared less interconnected and more punctate, although it was also closely co-distributed with the myosin II head probe staining in the CR region. In late stage, highly contracted CRs, STED imaging suggested that the anillin pattern appeared to become more network-like (Figure 8Q-U).

### Actin filaments are not required for septin2 or anillin recruitment to early CR clusters

Our previous work (Henson et al., 2017) and an earlier study from other investigators (Foe and van Dassow, 2008) have shown that treatment of sea urchin embryos with the actin filament disrupting drugs Latrunculin A or B does not disrupt the ability of myosin II to form clusters in the CR region during telophase. Latrunculin treated embryos do not divide and do not display the linearized pattern of aligned myosin II seen in control embryos (Henson et al., 2017). In order to test the actin dependence of septin2 or anillin localization to CR clusters we treated embryos with either LatA or LatB and then fixed and stained embryos and isolated cortices for P-MyoRLC and either septin2 or anillin at time points consistent with the timing of first division telophase in matched control embryos. We also monitored karyokinesis in these Lat treated embryos as this process is not inhibited by actin filament disruption. In both whole embryos (Figure 10A-C,G-I) and isolated cortices (Figure 10D-F,J-L) we saw evidence of an association between P-MyoRLC and septin2 (Figure 10A-F) or anillin (Figure 10G-L) in band-like arrays of clusters reminiscent of the early CR (Figures 4-6). Localization of nuclei in whole embryos indicated that they had undergone nuclear division (Figure 10C,I).

**Figure 10.**
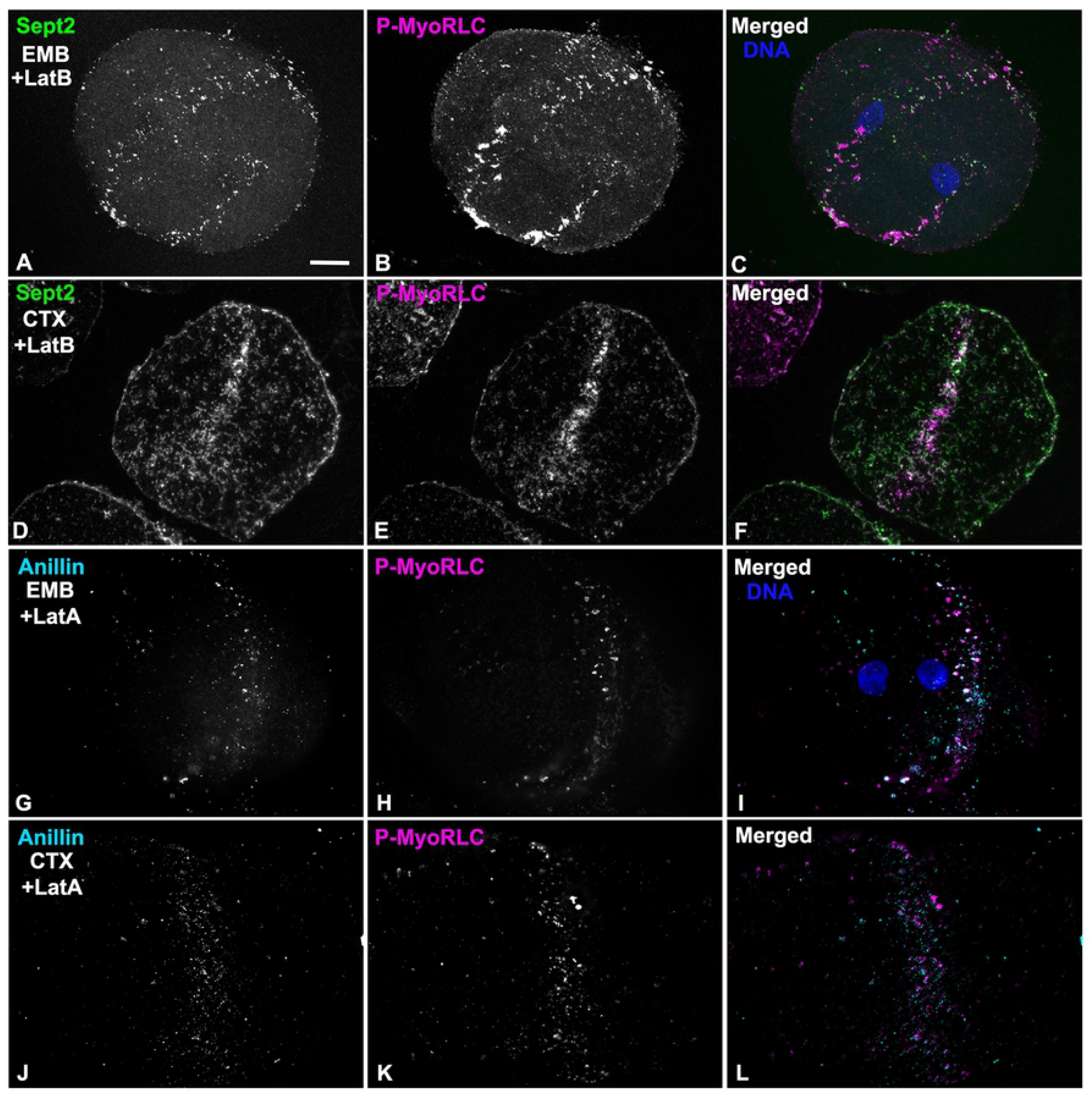
Actin filaments are not required for the recruitment of myosin II, septin2 or anillin to early stage CR clusters. (A-C) Whole embryo (EMB) treated with LatB and stained for septin2 (green), P-MyoRLC (magenta), and DNA (blue) and viewed using a through focus projection of a confocal Z series. The embryo contains a circumferential band of clusters containing myosin II and septin2 and the DNA staining indicates that it has undergone nuclear division during mitosis but not cytokinesis. (D-F) Cortex (CTX) isolated from a LatB treated embryo shows that septin (green) and P-MyoRLC (magenta) localize to a concentrated stripe of clusters. (G-I) Whole embryo (EMB) treated with LatA and stained for anillin (cyan) and P-MyoRLC (magenta) shows they colocalize in a stripe in an embryo which has undergone karyokinesis (I, DNA in blue). (J-L) Cortex (CTX) isolated from a LatA-treated embryo demonstrating a band of anillin and P-MyoRLC clusters. Bar = 10 µm in A; magnifications of A-L are equivalent. *L. pictus* embryos in A-F, *S. purpuratus* embryos in G-L.

## Discussion

Since the initial demonstration of the existence of the cytokinetic CR in the early 1970s by Schroeder (1970, 1972), there have been long standing questions about its precise architecture, dynamics, and regulation. As Glotzer (2017) pointed out in a recent review on cytokinesis, the structural organization of the metazoan CR remains poorly understood due to the rapid rearrangements and dense collections of filaments involved which make it difficult to resolve using conventional light microscopy. In our previous work (Henson et al., 2017) we have addressed these limitations by leveraging the advantage of the sea urchin embryo isolated cortex experimental system combined with super-resolution light microscopy and platinum replica TEM. Our results suggested that the CR assembles from an equatorial band of myosin II clusters that coalesce into a linearized arrangement of end-to-end myosin II filaments in the mature CR. Note that we are confident that the cortex isolation process itself does not induce these myosin II clusters as our imaging of whole embryos (Figure 3) and previously published work (Foe and von Dassow, 2008) have also indicated that clusters of myosin II are present in the CR. In the present study, we confirm and extend our recent work by once again using isolated cortices combined with super-resolution microscopy to explore the nanostructural organization and dynamics of the major CR scaffolding proteins anillin and septin.

### Anillin localization in the embryo CR highlights its potential scaffold functions

Since its initial discovery in *Drosophila* embryos (Field and Alberts, 1995), anillin is a protein that has been shown to localize to the animal cell cleavage furrow (Oegema et al., 2000; Maddox et al., 2005; Straight et al., 2005; Zhao and Fang, 2005), and functional studies have demonstrated that anillin is required for cytokinesis and serves as an essential scaffolding protein that links RhoA, the membrane, actin, myosin II, formin and septins (Oegema et al., 2000; Field et al., 2005; Maddox et al., 2005; Straight et al., 2005; Maddox et al., 2007; Piekny and Glotzer, 2008; Liu et al., 2012; Renshaw et al., 2014). Despite a great deal of attention given to the structure and function of anillin (Piekny and Maddox, 2010), the precise localization of anillin within the CR is not clear in animal cells, although it is well-defined in fission yeast. In these cells, the anillin-like protein Mid1 is localized in the nucleus, but upon entry into mitosis, it relocates to the cell surface around the nucleus, where it forms a number of small nodes (Bahler et al., 1998). Mid1 recruits the CR components myosin II (Myo2) and the formin Cdc12 (Laplante et al., 2016; Wu et al., 2006). Actin filaments nucleated from the nodes via formin activity are then captured by myosin II that in turn pulls the nodes together into a CR-like structure (Laporte et al., 2011; Laplante et al., 2016; Glotzer, 2017).

In the present study, we investigated anillin localization in the sea urchin embryo CR by first characterizing the specificity of an antibody raised against the PH domain of sea urchin anillin by confirming its appropriate immunoreactivity in immunoblotting and predicted localization in the nuclei of interphase cells (Figure 1). In whole embryos we show that sea urchin anillin is associated with the cleavage furrow in the expected orientation relative to the mitotic apparatus (Figure 2) and that it colocalizes with the CR marker activated myosin II, P-MyoRLC (Figure 3). In cortices isolated from first division embryos, conventional imaging revealed that anillin associates with myosin II initially in clusters/nodes that then progress to narrower, more linearized arrays in more mature CRs (Figure 4; 5). Super-resolution imaging emphasized the close association between anillin and myosin II within the early stage clusters (Figure 6; 7), with anillin often displaying a C or O-shaped structure in cluster centers, as well as the possibility suggested from Z axial images that anillin resides closer to the membrane relative to myosin II head groups. The fact that anillin localizes to the CR region in clusters independent of actin filaments (Figure 10) is consistent with previous work in *Drosophila* S2 cells (Hickson and O’Farrel, 2008), but inconsistent with results from BHK-21 mammalian cells (Oegema et al., 2000).

Our anillin localization results are in general agreement with previous fPALM super-resolution imaging of fission yeast nodes (Laplante et al., 2016) and mature CRs (McDonald et al., 2017) which suggest that the anillin-like Mid1 resides in the center of nodes and nearest the membrane relative to other node/CR proteins. Higher order anillin structures in sea urchin CRs are suggested by 3D-SIM images of late stage CRs where anillin on occasion appears reticular (Figure 8). Earlier work has hinted at organized staining patterns for anillin in the CR, to include the punctate distribution of anillin along actin cables in the cleavage furrow of BHK cells (Oegema et al., 2000), the presence of patches or filaments containing anillin and myosin II in the CR of HeLa cells either untreated or arrested in cytokinesis by blebbistatin treatment (Straight et al., 2003; 2005), and the presence of combined anillin and septin rings in the intercellular bridge region of late stage dividing HeLa cells (Renshaw et al., 2014).

It is important to note that the predicted multi-domain nature of sea urchin anillin (Figure 1A) is supported by our ability to colocalize this protein with structures in the CR containing its expected binding partners – namely myosin II, actin, septin, and RhoA (Figure 3-8). This localization pattern provides strong circumstantial evidence that anillin is playing its expected role as an integrating scaffold in the sea urchin CR. We speculate that anillin recruitment via RhoA activated from the astral microtubule trafficked centralspindlin complex (Argiros et al., 2012) is the initial event in the establishment of the clusters that initiate the process of CR formation.

### Septin2 localization in the sea urchin embryo CR demonstrates higher order structure

Septins are a family of G proteins first discovered in yeast (Hartwell, 1971) that can combine into hetero-oligomeric filaments which consist of different isoforms in different species, and which play important roles in cell polarity, membrane remodeling, morphogenesis, exocytosis, and cytokinesis (Bridges and Gladfelter, 2015; Marquardt et al., 2019). Septins have long known to be associated with the CR in many different cell types (Bridges and Gladfelter, 2015) and septin2 in particular has been shown to bind to myosin II and facilitate the myosin II activation required for cytokinesis (Joo et al., 2007). In the present study, we demonstrate that septin2 associates with myosin II, actin and anillin in clusters that appear to serve as the antecedent of the CR (Figure 4-9). Within these clusters septin2 is more central than myosin II (Figure 6; 7), in the same area as anillin, with XZ imaging suggesting little axial separation between septin2 and myosin II. In fission yeast CR precursor nodes do not contain septin (Glotzer, 2017), although septin filaments are present in the mature CR (McDonald et al., 2017) and septin filaments form characteristic hourglass and double ring structures in the CR-equivalent structure in budding yeast (Ong et al., 2014). Our 3D-SIM and STED images indicate the presence of elongate septin2 filaments associated with enlarging clusters and with the mature CR (Figure 6-9). These septin2 filaments associate with but do not precisely colocalize with myosin II staining and frequently display periodic labeling, as might be expected if the sept2 antibody is labeling just one septin isoform within a hetero-oligomer. In some images the septin2 staining is elongate (Figure 6; 8; 9) which may represent the ability of short septin filaments to diffuse in the membrane and anneal into longer filaments via end-to-end associations as has been demonstrated *in vitro* on phospholipid bilayers and *in vivo* in fungi (Bridges et al., 2014). The ability of septin2 to localize to the CR region independent of actin filaments (Figure 10) is similar to the results reported for the septin Pnut in LatA-treated *Drosophila* S2 cells (Hickson and O’Farrell, 2008).

The higher order structural organization of septin2 filaments present in mature stage CRs of sea urchin embryos appears to be a reticular network with an overall honeycomb pattern which is closely associated with the distribution of myosin II (Figure 8; 9). The imaging does not allow for a distinction between an interconnected network of filaments versus an array of separate filaments in an overlapping organization. However, this septin2 network is similar to the gauze-like septin filament structures that have been previously reported *in vitro* using negative stain TEM (Garcia et al., 2011; Bertin et al., 2010) and *in vivo* associated with the cortex of budding yeast using platinum replica TEM (Rodal et al., 2005) and TEM tomography (Bertin et al., 2012), and associated with the interface of transverse and dorsal stress fibers in the leading lamella of migrating mammalian epithelial cells using SIM (Dolat et al., 2014). Prior work with dividing mammalian tissue culture cells has shown filamentous septin staining codistributed with CR actin cables (Oegema et al., 2000) and septin rings in the midbody at the end of cytokinesis (Renshaw et al., 2014). However, we believe that our super-resolution images provide the first visualization of a clearly defined septin filament higher order organization in the CR of an animal cell.

### Clusters of myosin II, anillin and septin initiate the sea urchin embryo CR

Our results suggest that an equatorial band of cortical clusters of myosin II, septin2 and anillin help serve as a precursor for the formation of the CR in first division sea urchin embryos. Earlier work in *C. elegans* embryos (Maddox et al., 2005; 2008) and in *Drosophila* S2 cultured cells (Hickson and O’Farrell, 2008) have indicated that clusters of these same three proteins are also involved in the cytokinesis process, although these clusters are not completely analogous to our results and the imaging employed is significantly lower resolution. In *C. elegans* the anillin homologue ANI-1 is required for the cortical ruffling and pseudocleavage that proceed cytokinesis as well as for asymmetric cleavage patterns during cytokinesis, although CR closure can occur in the absence of ANI-1 (Maddox et al., 2005; 2007). In addition, ANI-1 targets independently to the CR and helps direct septins, but not myosin II, there as well. In contrast with the equatorial band of clusters we demonstrate in the sea urchin embryo, the clusters in *C. elegans* appear distributed throughout the entire cortex of the embryo during ruffling, although clusters of myosin II do appear in the CR region during cytokinesis (Osório et al., 2019). In S2 cells the CR localization of anillin is independent of actin and myosin II, its absence causes destabilization of the position of the cleavage furrow, and aggregates of myosin II, anillin and septin in the CR only become obvious upon treatment of dividing cells with Latrunculin (Hickson and O’Farrell, 2008). These aggregates can appear filamentous with myosin II and anillin staining appearing in the same place but slightly offset (Hickson and O’Farrell, 2008), reminiscent of what we demonstrate in our SIM images of anillin and myosin II staining of clusters. In general, it appears that the cytokinetic myosin II, anillin and septin clusters in *C. elegans* embryos and in *Drosophila* S2 cells are loosely analogous with those we demonstrate in the sea urchin embryo. However, the differences outlined above, the superior spatial resolution of the architecture of clusters we provide, as well as the strong evidence of the transformation of the sea urchin clusters into patches and linearized arrays makes it difficult to make more direct comparisons.

### A working conceptual model for how the CR is built in the early sea urchin embryo

Our current conceptual model of CR formation in the first division sea urchin embryo employs the following hypothetical framework as informed by the results of our present study. At late anaphase the astral microtubule-dependent activation of Rho via the action of centralspindlin/RhoGEF helps trigger the recruitment of anillin, myosin II and septin to the CR precursor clusters (Figures 4-7), which is facilitated by the ability of these three CR components to bind to one another. Within the clusters anillin and septin occupy a central core (Figures 6; 7) and would be expected to interact with the membrane and bind with the tails of myosin II filaments. Actin filaments are not necessary for the recruitment of myosin II (Foe and von Dassow, 2008; Henson et al., 2017), septin (Figure 10) or anillin (Figure 10) to the CR region in sea urchin embryos and research in other cell types suggests that anillin recruitment is independent of myosin II, whereas septin recruitment is dependent on anillin (Maddox et al., 2005; 2007; Hickson and O’Farrell, 2008). As noted earlier, the sea urchin clusters share some gross similarities with the organization of clusters of these proteins in dividing *C. elegans* embryos and *Drosophila* S2 cells as well as the nodes seen in fission yeast undergoing cytokinesis. However, the yeast nodes do not contain septin and the orientation of the myosin II filaments in our images suggests a mini-filament chain organization instead of the bouquets of individual myosin II proteins defined by fPALM imaging of yeast nodes (Laplante et al., 2016). Over time the sea urchin clusters enlarge with myosin II forming head-to-head filament arrays oriented parallel to the plane of the membrane via interaction with actin filaments, septin filaments elongating via annealing with other septin oligomers (Bridges et al., 2014), and the anillin array also expanding (Figure 6; 7). In fission yeast node-associated formins nucleate actin filaments which then are used by node-associated myosin II to move nodes towards the forming CR (Valvylonis et al., 2008). Actin filaments do associate with the sea urchin clusters (Figure 5), however the distribution of formin has not been established in our system. Coincident with cluster enlargement these structures tend to interconnect and coalesce into a narrower and denser linear array in which the interaction of myosin II with elongate actin filaments causes them to become aligned parallel with the axis of the cleavage furrow (Figure 5; 8), septin filaments organize into a network (Figure 8; 9), whereas anillin tends to remain in more punctate arrays until late in cleavage where they also appear to form a network (Figure 8).

Within the context of the mature CR, septin and anillin can be considered to be functioning as scaffold and anchoring proteins although they also may engage in other critical roles. For example, septins may be contributing to the full activation of myosin II by serving as a platform for myosin II and its activating kinases (Joo et al., 2007), and/or they may be essential for the curvature and bundling of the actin filaments in the CR (Mavrakis et al., 2014; Bridges and Gladfelter, 2015). In the sea urchin embryo septin and anillin remain in the CR through the midbody stage (Figure 2, 3) and therefore may be expected to play a role in the maturation of the CR to the midbody ring and subsequent abscission given the evidence that they participate in this critical step in other cells (Etsey et al., 2010; Kechad et al., 2012; El Amine et al., 2013; Renshaw et al., 2014; Karasmanis et al., 2019). In a recent interesting hypothesis paper, Carim et al. (2020) argue that anillin is critical for the organization of two Rho-dependent sub-networks during cytokinesis: the acto-myosin sub-network driven by functionality associated with anillin’s N terminus and the anillo-septin sub-network dependent on anillin’s C terminus. The anillin localization patterns that we demonstrate in the present study would appear to be consistent with this potential central organizing role for anillin.

In conclusion, the results of the present study suggest that CR generation in sea urchin embryos may be an evolutionary derivative of the process in fission yeast, with both mechanisms relying on initiation via clusters/nodes of crucial cytokinetic proteins, including myosin II, anillin, actin, and, in the sea urchin, septin. In order to further explore this evolutionary relationship and to address the many questions raised by our conceptual model, we are currently pursuing a number of new experimental directions. These include CR imaging approaches in live embryos to more closely examine dynamics, investigations into the role of formins in CR formation, and development of agent-based computer modeling of cytokinesis informed by our microscopy-based studies.

## Acknowledgements

Grateful thanks are extended to Dr. Simon Watkins and Michael Calderon (Center for Biologic Imaging, University of Pittsburg) for their expert assistance with STED microscopy, to Dr. Billie Swalla (University of Washington) for access to instrumentation and reagents at Friday Harbor Laboratories (FHL), and to Casey Ditzler, Aphnie Germain, Patrick Irwin, Eric Vogt, Erik Williams, Bakary Samasa, and Ethan Burg (Dickinson College) for help with experimentation and image analysis. This research was supported by National Science Foundation (www.nsf.gov) collaborative research grants to J. H. H. (MCB-1917976) and C. B. S. (MCB-1917983). C. G. was supported by the Charles Lambert Fellowship and the FHL Research Fellowship Endowment.

## Supporting Information

**S1 Spreadsheet. Data sets used for graphs in Figs 4T, 6N, 6O and 7L**.

